# Low-Density EEG for Neural Activity Reconstruction Using Multivariate Empirical Mode Decomposition

**DOI:** 10.1101/713610

**Authors:** Andrés Felipe Soler, Pablo A. Muñoz-Gutiérrez, Maximiliano Bueno-López, Eduardo Giraldo, Marta Molinas

## Abstract

Several approaches can be used for estimating neural activity. The main differences between them are in the apriori information used and their sensibility to high noise levels. Empirical Mode Decomposition (EMD) has been recently applied to Electroencephalography EEG-based neural activity reconstruction to provide apriori time-frequency information to improve the neural activity estimation. EMD has the specific ability to identify independent oscillatory modes in non-stationary signals with multiple oscillatory components. The various attempts to use EMD in EEG analysis, however, did not provide yet the best reconstructions due to the intrinsic mode mixing problem of EMD. Some previous works have used a single-channel analysis and in other cases, multiple-channel have been used for other applications. In this paper, we present a study about multiple-channel analysis using Multivariate Empirical Mode Decomposition (MEMD) as a method to attenuate the mode mixing problem and to provide apriori useful time-frequency information to the reconstruction of neuronal activity using several low-density EEG electrode montages. The methods were evaluated over real and synthetic EEG data, in which the reconstructions were performed using multiple sparse priors (MSP) method with several electrode numbers of 32, 16, and 8, and the source reconstruction quality was measured using the Wasserstein Metric. Comparing the solutions when no pre-processing was made and when MEMD was applied, the source reconstructions were improved using MEMD as apriori information in the low-density montage of 8 and 16 electrodes. The mean source reconstruction error on a real EEG dataset was reduced a 59.42% and 66.04% for the 8 and 16 electrodes montages respectively, and on a simulated EEG with three active sources, the mean error was reduced an 87.31% and 31.45% for the same electrodes montages.

## I. Introduction

EEG is an indicator of neural activity and it is used to study complex brain dynamic processes, as the cognitive processes, memory process, and emotion recognition Soleymani et al. (2016), Lin et al. (2016). Due to the non-linear and non-stationary nature of EEG signals, their analysis is a challenging task in either, time and frequency domains. However, some hidden information can be extracted to be used in early detection of different disorders by using advanced techniques in signal processing and analysis Subha et al. (2010). In recent years, Hilbert Huang Transform (HHT) is increasingly used in the analysis of such signals Bueno-López et al. (2018). However, in some applications, the extraction of information has been hampered by the mode mixing problem that appears in the Empirical Mode Decomposition (EMD) when frequency components are relatively close or exhibit intermittency. A mode mixing problem can be identified when a set of signals of widely disparate scales appear in an Intrinsic Mode Function (IMF), or when a signal with a similar scale appears in different IMF components. Mode mixing is a consequence of spectral proximity of the frequency components or signals intermittency.

The presence of mode mixing can hamper the physical interpretation of the process which is intended to be described by the individual IMFs Wu and Huang (2009), Rilling and Flandrin (2008). The mode mixing problem has been studied in applications in several fields, for example in Xue et al. (2016), the authors discuss the mode mixing influence on the hydrocarbon detection based on EMD. They use several variations of EMD to eliminate the mode mixing effects, specifically, they apply Ensemble EMD (EEMD) and Complete Ensemble EMD (CEEMD), as useful tools to identify the peak frequency volume and the peak amplitude above-average volume.

In 2009, a new strategy was presented in Rehman and Mandic (2009). In which, a multivariate variation of EMD called Multivariate Empirical Mode Decomposition (MEMD). The MEMD is a method that reduces the mode mixing problem and it is a good alternative for multichannel data analysis, as is the case of EEG signals. Some previous papers have reported the use of EMD for neural activity reconstruction and other applications in bioengineering Bueno-Lopez et al. (2017a), Bueno-Lopez et al. (2017b), Okcana (2016), Men-Tzung et al. (2008), but the use of MEMD for the same application is less. In Yin et al. (2012) the authors presented a method for data analysis based on MEMD, applying a pre-processing step with Independent Component Analysis (ICA) to calculate and evaluate the energy presented in an EEG record from the quasi brain deaths and to evaluate the brain activity. In Zahra et al. (2017), the authors proposed a data-driven method for classifying ictal (epileptic activity) and non-ictal EEG signals using MEMD algorithm. In which, they extract and select suitable feature sets to classify the neural activity based on a multiscale Time-Frequency representation of the EEG signals by the application of MEMD. A fusion between MEMD and source reconstruction algorithms with an unsupervised eye blink artifact remover was introduced in Khosropanah et al. (2018), with the purpose of accurate localization of epileptogenic sources. In She et al. (2017) a novel identification method of relevant Intrinsic Mode Functions is proposed based on Noise-assisted-MEMD and Jensen-Shannon distance measure.

In this research work, we propose the application of multivariate time-frequency EEG signals analysis using the MEMD method as a pre-processing step before applying source localization algorithms. The MEMD decomposes the signal in several intrinsical mode functions IMFs, in which the information of the underlying brain activity is separated in frequency bands. Due to the relation between the information in each channel, MEMD reduces the mode mixing problem, which allows understanding the effect of a stimulus on different regions of the brain. Next, selected information of the decomposed EEG signals are used to perform neural activity reconstruction with high accuracy than using the raw electrode information, therefore, less number of electrodes is required to extract this underlaying time-frequency information due to the MEMD properties and the EEG redundancy in the electrode measurements on the scalp. This hypothesis is evaluated with simulated EEG data and real EEG signals from a face-evoked potentials paradigm.

We perform MEMD on the sets of EEG data and then perform source activity reconstruction using Multiple Sparse Priors (MSP). According to Jatoi and Kamel (2018) MSP method presents better performance than other well-known algorithms to solve the EEG inverse problem like Minimum Norm Estimation (MNE), Low-Resolution Tomography (LORETA), and beamforming, using a reduced set of seven electrodes. In our case, we evaluated the source reconstruction with 32, 16 and 8 electrodes to analyze and discuss the effects of channel reduction with the proposed methodology.

## II. Material and Methods

### A. Empirical Mode Decomposition

The Empirical Mode Decomposition (EMD) is a method adaptive, which dependent directly from data. The aim this method is to decompose a nonlinear and non-stationary signal ***y***(*t*_*k*_) into a several intrinsic mode functions (IMFs), in which each one satisfies the two following conditions Huang et al. (1998):

1. The number of extrema and the number of zero crossings must be the same or differ at most by one.
2. At any point, the mean value of the envelope defined by the local maxima and the envelope defined by the local minima is zero.

Empirical Mode Decomposition is applied over ***y***(*t*_*k*_) to obtain ***γ***_*i*_(*t*_*k*_) being *i* the intrinsic mode function (IMF), and

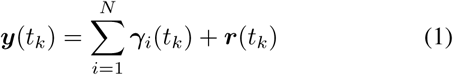

where *N* represents the number of IMFs and ***r***(*t*_*k*_) a residual information. Recently, some optimization techniques have been proposed to improve the performance of the EMD Xu et al. (2016), Hou and Shi (2013).

Having obtained the intrinsic mode function components, we can apply the Hilbert transform to each IMF component, and compute the instantaneous frequency according to equation (2).

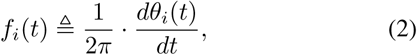

where *θ*_*i*_(*t*) is the function phase of each IMF calculated from the analytic signal associated Boashash (1992). Finally, the instantaneous frequency can be observed in the Hilbert Spectrum.

### B. Multivariate Empirical Mode Decomposition (MEMD)

The MEMD allows a decomposition of the EEG ***y***(*t*_*k*_) in several IMFs ***γ***(*t*_*k*_) and a residual ***r***(*t*_*k*_) as follows:

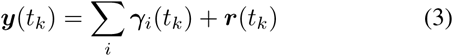

The brain mapping can be performed either for each ***γ***_*i*_(*t*_*k*_) independently or for a linear combination of IMFs.

### C. IMF selection: Entropy Function

An entropy based cost function is applied over each IMF ***γ***_*i*_(*t*_*k*_) as follows (Bueno-López et al. (2019), Muñoz-Gutiérrez et al. (2019)):

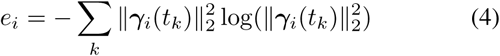

being *e*_*i*_ the entropy of each IMF, and ***e*** = [*e*_1_ … *e*_*N*_]. With the objective to rebuild the EEG signal 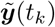 a selection of the IMFs, based on the IMFs with highest entropy, is applied according to the measured entropy *e*_*i*_.

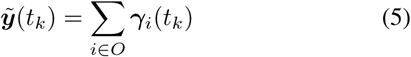

in which *O* represents the subset of IMFs that have been selected to build the filtered EEG signals.

### D. Source Reconstruction Algorithm

The relation between the brain activity at cortical areas (source activity) and the measurement electrodes in the scalp is represented by the forward problem equation:

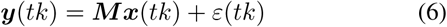

being ***y***(*tk*) ∈ ℝ^*d×T*^ the EEG signals at *d* number of electrodes with *T* samples, ***x***(*tk*) ∈ ℝ^*n×T*^ contains the amplitude of *n* sources distributed over the cortical areas, *ε*(*tk*) is a noise covariance, assumed as a Gaussian distributed with zero mean, and ***M*** ∈ ℝ^*d×n*^ is the lead field matrix or volume conductor model, and it represents the physical model based on head anatomy, and explains how the potentials flow through from brain to electrodes, this is based on the conductivities of different layers between current sources and electrodes, like scalp, skull, CSF, gray matter, and white matter.

The source reconstruction involve the solution of the EEG inverse problem, which is mathematically ill-posed and ill-conditioned, due to the high number of unknowns sources of activity (thousands of sources), and the reduced quantity of observations (tens of electrodes). To overcome such challenging characteristics, several approaches tackle the inverse problem as a minimization problem with spatial constraints, in which, the solutions are smooth in the source space as MNE (Hämäläinen and Ilmoniemi (1994)) and LORETA (Pascual-Marqui et al. (1994)). In other hand, the MSP method with a Bayesian approach has been gaining attention due to the sparse solutions outcome and the high performance in terms of localization error and free energy, as presented in Friston et al. (2008) and validated in López et al. (2014) and Jatoi and Kamel (2018). Due to outcomes of MSP, we chose it to perform the brain mapping and to evaluate the effects of using MEMD as a prior information. This method can be found in the SPM12 software (Friston et al. (2007)) as a package for MATLAB (The MathWorks, Inc.)

### E. Generation of synthetic EEG Signals

To assess the solution for the neuromagnetic inverse problem using EEG signals, we performed simulations with several scenarios in which the brain activity is known. Therefore, it is necessary to use a lead field matrix (head model, preferably a realistic representation) that allows generating EEG signals with active sources in predefined or random positions and with specific activity function.

The head model used to generate the synthetic EEG signals, can be found it in the database called *Multi-modal Face Dataset* in http://www.fil.ion.ucl.ac.uk/spm/data/mmfaces/ of SPM software. This dataset was obtained using the same paradigm reported in Henson et al. (2003) and contains EEG, MEG and fMRI data for one subject. This paradigm has been used in different works for source reconstruction and addressed to evoked responses (Friston et al. (2006); Henson et al. (2007, 2009, 2011); Gramfort et al. (2013); Fukushima et al. (2015)). The head model contains a cortical mesh with 8196 vertices as distributed sources and relates them to 128 electrodes. However, reduction to 8, 16 and 32 channels is performed as shown in Fig. 1. This reduction in the number of measurements is performed in order to analyzed the quality of the reconstruction vs the number of measurements.

**Fig. 1:**
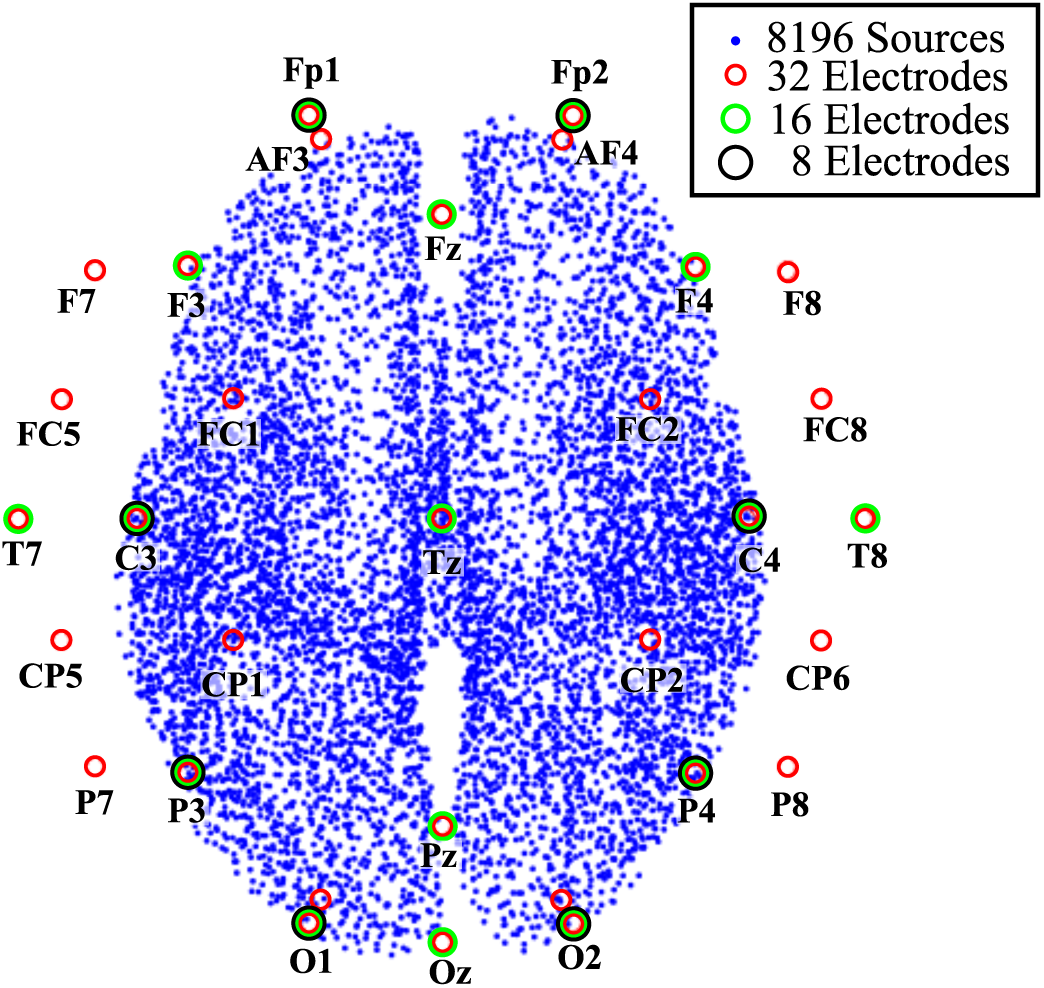
Three configurations of electrode positions for EEG measurements.

Nine configurations of EEG signals were proposed. We considered three different number of active sources: 1, 3, and 5. For each number of active sources, the synthetic EEG was generated considering 8, 16 and 32 electrodes. One of the goals, for this paper, was to show that the MEMD was able to reduce the mode mixing generated in the process to obtain the IMFs. MEMD allows to make a better choice of frequencies inside the each IMFs, therefore, the source activity for 1, 3 and 5 active sources were simulated with different frequencies and different instants of time. Moreover, the evaluation was done in different levels of measurement resolution, i.e. 8, 16 and 32 channels as it is shown in Fig. 1.

The brain activity was simulated for each source using a windowed sinusoidal activity, as follows:

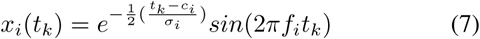

The signals were generated in a time interval from 0s to 6s with a sample frequency *f*_*s*_ of 200*Hz*. The activities frequencies were selected into the typical ranges for brain frequencies, e.g., *alpha* or *beta* brainwaves, in order to have a realistic scenario, in addition, in the cases of 3 and 5 active sources, some sources were located at low brain frequencies e.g., *theta* or *delta*, to observe the perform under several range of frequencies. For the case of 1 active source, the activity was generated in *t* = 1*s* with a *f* = 10*Hz* and the active source was in the position 4000, Fig. 2a shows the configuration for this first case

**Fig. 2:**
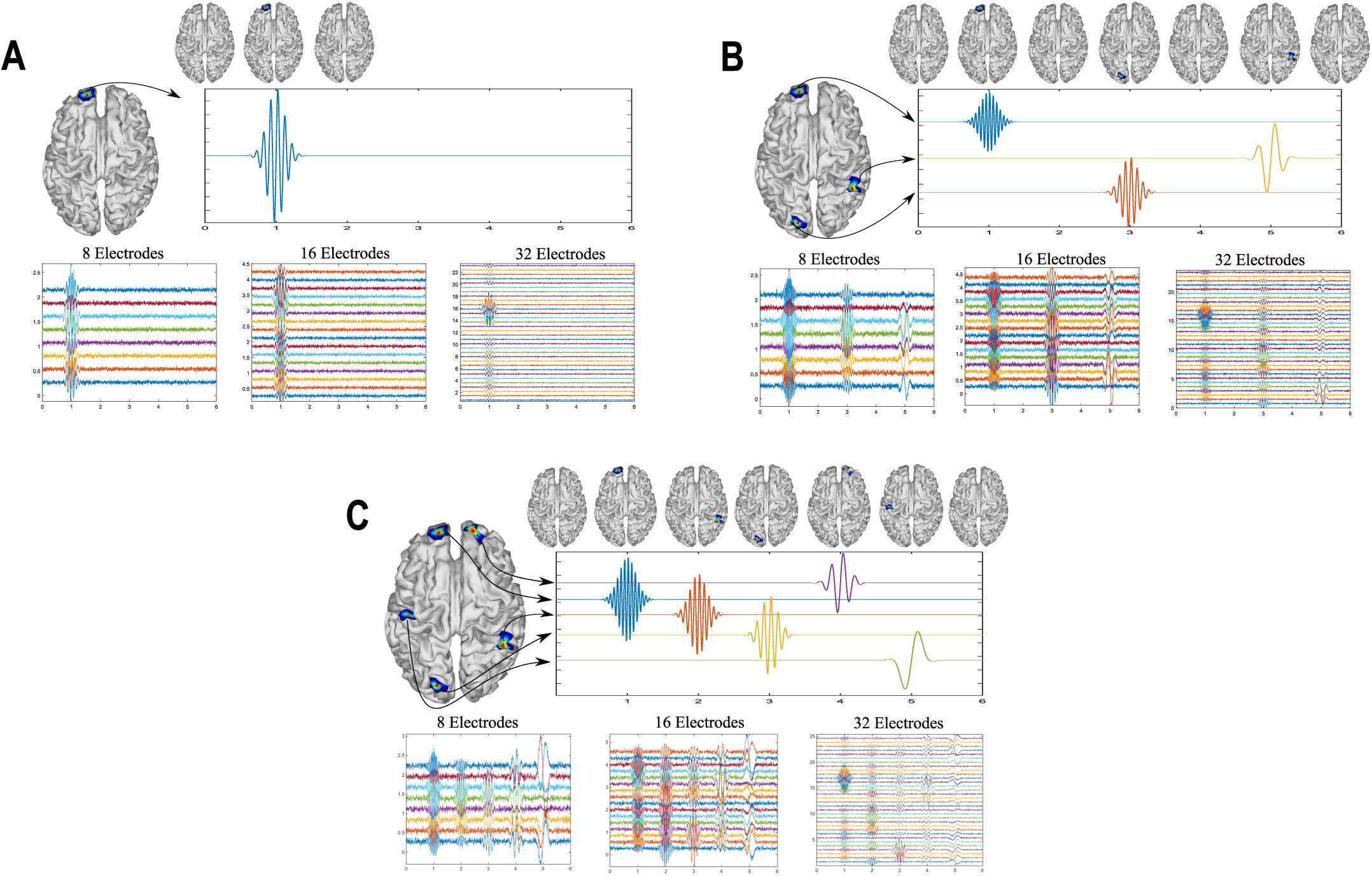
Simulated activity for one source **(A)** with 10Hz and 32, 16 and 8 EEG channels. Simulated activity for three sources **(B)** with 4, 12 and 20Hz windowed sinusoidal activity and using 32, 16 and 8 EEG channels. Simulated activity for five sources **(C)**with 2, 6, 10, 15 and 20Hz windowed sinusoidal activity and using 32, 16 and 8 EEG channels

In the cases of 3 and 5 active sources, the activity were simulated in different instants of time. In Fig. 2b are shown the simulated signals for 3 active sources. For the case of 3 active sources, the first activity was generated in the source located in the position 4000 at *t* = 1*s* with a *f* = 20*Hz*, at *t* = 3*s* with a *f* = 12*Hz* was activated the second source in the position 5020 and, the third source, was located in the position 150 at *t* = 5*s* with *f* = 4*Hz*.

Finally, for 5 active sources the signals were generated at *t* = 1*s, t* = 2*s, t* = 3*s, t* = 4*s* and *t* = 5*s*, with *f* = 20*Hz* in the position 4000, with *f* = 15*Hz* in the position 5020, with *f* = 10*Hz* in the position 150, with *f* = 6*Hz* in the position 8100 and with *f* = 2*Hz* in the position 2200, respectively. Fig. 2c shows the simulated activity for 5 sources.

A common evaluation of inverse problem solutions is performed by using simulated sources where the underlying activity is known. In this work, simultaneous simulated sources with sinusoidal activity are used in order to evaluate the performance of the MEMD in terms of the separability of the source activity in the frequency domain. To this end, simulated EEG activity is obtained for 1, 3 and 5 sources located spatially and temporally at different points. Sources frequencies *f*_*i*_ in the range of 2Hz to 20Hz are considered. Also, temporal localization of sources is in the range of 1 to 5 seconds. The source reconstruction is made in two ways, using MSP directly from the synthetic EEG without any pre-processing step, like raw signals, and using MEMD as prior information to MSP, in which the main IMFs were selected according to the entropy function in equation 4, where those IMFs whose presented the highest entropy were used to recalculate the EEG.

The simulating procedure starts with the generation of each source, using the windowed sinusoidal activity, the active sources are located in predetermined locations, the activity ***x***(*t*_*k*_) is calculated, and then, the synthetic EEG is obtained by using:

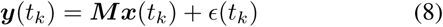

Then a noise is added to the EEG signal ***y***(*t*_*k*_) with a signal-to-noise ratio of *SRN* = 10. Where three configurations are considered for the measurements: 32, 16 and 8 EEG channels. For each synthetic EEG at different channel number, a reduced lead field matrix were used to perform the brain mapping and then, the source reconstruction were calculated directly with MSP (raw MSP) and applying MEMD to the electrode space (MEMD-MSP). Finally the reconstruction were compared with a spatial accuracy measurement, the Wasserstein Metric.

### F. Real EEG Signals Database

For an evaluation of the MEMD method and his application over real EEG signals, a multi-subject, multi-modal human neuroimaging dataset was used. The experiment has 16 participants where the stimulus were presented in a screen to each one of the subjects, three conditions were considered in the stimulus, familiar faces (famous), unfamiliar faces (non-famous) and scrambled faces. As described in Wakeman and Henson (2015), the experiment involved EEG, MEG and fMRI to estimate the neural activity and their location over the cortical areas of brain by applying a multi-modal technique presented in Henson et al. (2011). The EEG recordings were taken with 70 electrodes of AgCl using a layout according to the 10-10 system. All the subjects in database have their own head model and their ground truth activity. The lead field matrix that modeled the head conductivities was made by 8196 distributed sources. The lead field matrix was used for solving the inverse problem and the ground truth activity was used to compare the obtained solutions with the MEMD method as apriori information to MSP and the EEG directly with MSP. As the dataset contains the ERPs of the experiment, we considered the event related potentials ERPs around the 170 ms or ‘N170’ component for the experiments with scrambled and familiar faces. As well like in the simulated data, a reduction of channels was developed from 70 electrodes to 32, 16 and 8, in order to evaluate the performance of the brain mapping solution with MSP using one or several IMFs from MEMD and comparing the results against MSP with raw data. For evaluate the activity around the N170, we have established the region of interest ROI as in Henson et al. (2011), in the window between 100 ms and 220 ms. The methodology followed for processing the high density data is presented in Fig. 3.

**Fig. 3:**
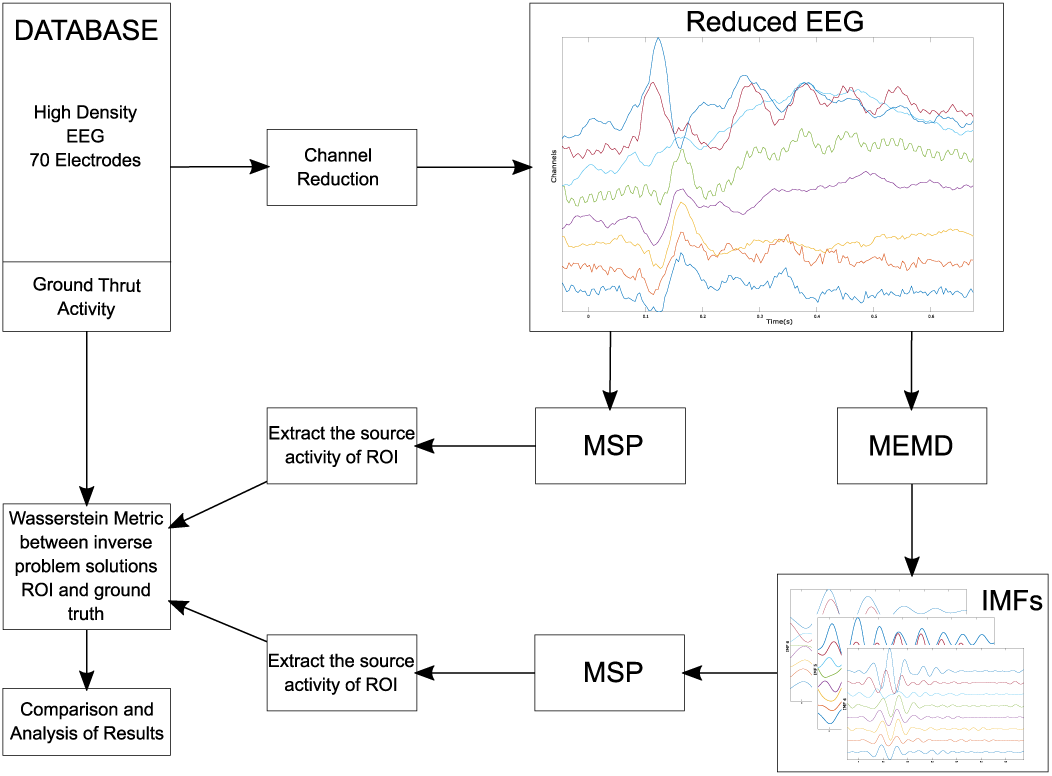
Block diagram of the methodology followed for processing the EEG from database.

According to the number of electrodes to evaluate, the channels are extracted from the high density EEG. The reduced data is directly processed by MSP to obtain the called raw inverse solution. Besides, the reduced data is also processed with MEMD and one or several IMFs are selected to obtain the inverse solution with MSP. Finally, the both reconstructions of the source activity are compared with the ground truth to measure the spatial accuracy of the solution using the Wasserstein Metric. Following a similar procedure as in the synthetic EEG signals, the lead field matrix was reduced according to the position of electrodes, and we selected the channels trying to maintain an equal spatial distribution of them over the scalp. The layout of the reduction is presented in Fig.4a, 4b and 4c for 32, 16 and 8 electrodes respectively. In addition, Fig.4d shows the 8196 distributed sources of one of the subjects, the reduction of electrodes for 32, 16, and 8 channels, and their positions.

**Fig. 4:**
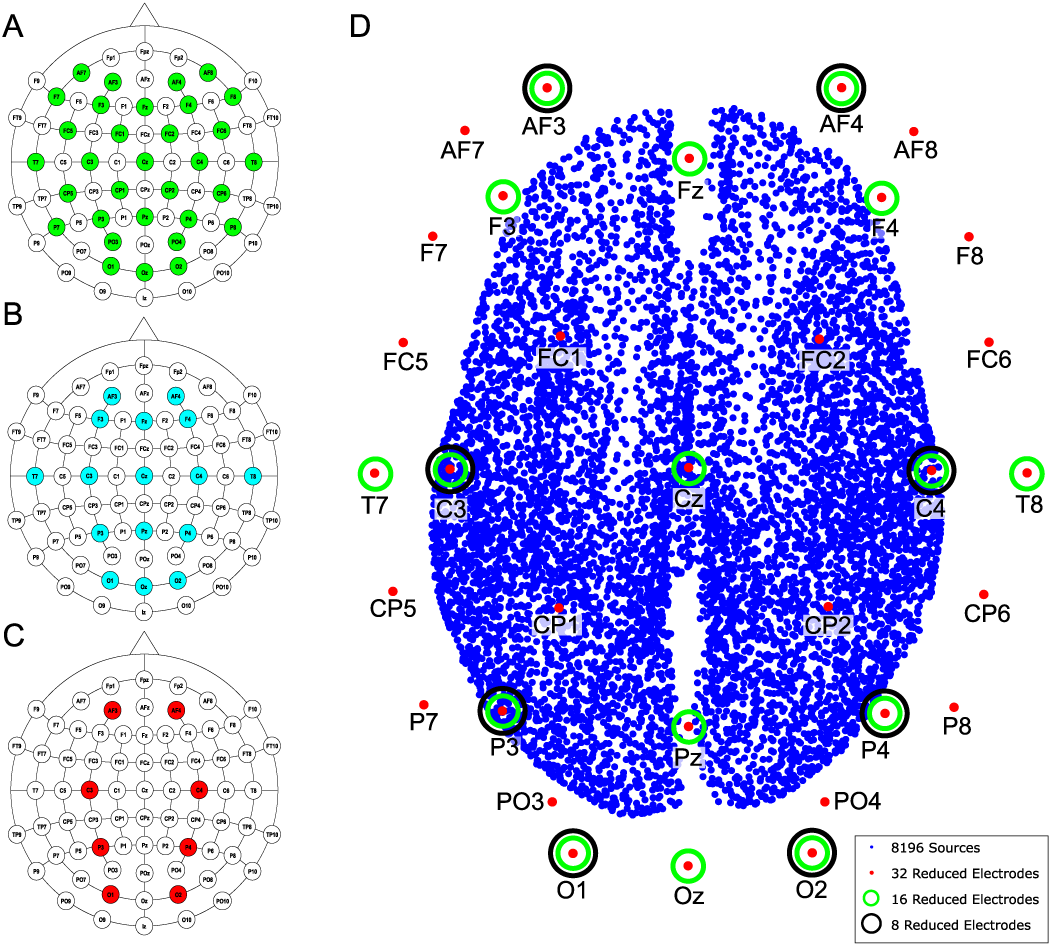
Layout according to the 10-10 system for 70 electrodes and the performed reduction for **(A)** 32 electrodes, **(B)** 16 electrodes, **(C)** 8 electrodes,**(D)** Brain model with 8196 distributed sources, names and position of electrodes used in the channel reduction.

### G. Accuracy Measurement

The Wasserstein metric *w*_*m*_ (also known as *Earth-Movers Distance*, Rubner et al. (2000)) is used as a quality index of the source reconstruction accuracy. This measured provide a spatial comparison between the ground truth and the estimated source activity, in which, the *w*_*m*_ ∈ ℝ^+^ measures the work to transform the estimated power distribution of sources into the ground truth power distribution by “transporting” probability mass. (Castano-Candamil et al. (2015)). With this index, a lower *w*_*m*_ value represents a better spatial accuracy of the source reconstruction. This metric have been used to compare EEG/MEG inverse solutions in order to have a meaningful measure for the estimated source distributions (Haufe et al. (2008)). As the Wasserstein metric is considered as a spatial accuracy index, usually the mean activity during the complete EEG segment is compared with the mean ground truth activity to assess the source localization. In addition, to offer a temporal assessment of the reconstructed activity, the solutions also are evaluated using small time windows (time-ROIs), in the case of synthetic EEG, the time-ROIs are defined 250 ms before and 250 ms after the time of maximum activity of each one of the simulated sources, and for the real dataset assessment, the ROI is definet between 100 ms and 22 0ms. In general, the mean activity in the time-ROI is calculated and then compared with the mean activity of the ground truth during the same time-ROI.

## III. Results

### A. Synthetic EEG Data Study

We generated and processed the EEG signals for the nine aforementioned cases. An example of the application of MEMD for the reconstruction case of one source with 10Hz and 8 EEG channels are shown in Fig. 5. It can be seen that MEMD shows in the IMF2 noise activity and there is not any identifiable source activity, meanwhile, the IMF3 unmixed the source activity, it is clearly identifiable with no underlying noise. In addition, when the brain activity reconstruction is obtained, it can be seen that for the MSP without preprocessing using MEMD (raw-MSP), the source activity is split in two sources, in which one of them has an acceptable location, however, the main activity represented with a red color is located in the prefrontal cortex but in a lateral position far of the original source, this fact explains the higher value of *w*_*m*_ = 6.23 than MEMD-MSP reconstruction. In other hand, by using the MEMD-MSP has obtained *w*_*m*_ = 1.26, and in the source map the main activity is accurately located, even if spurious activity appears at the same position than with raw MSP, its value is attenuated. The appearance of this “ghost” activity could be due to the performed channel reduction, however, is remarkable that the value with MEMD-MSP is lower and the main activity is clearly identifiable.

**Fig. 5:**
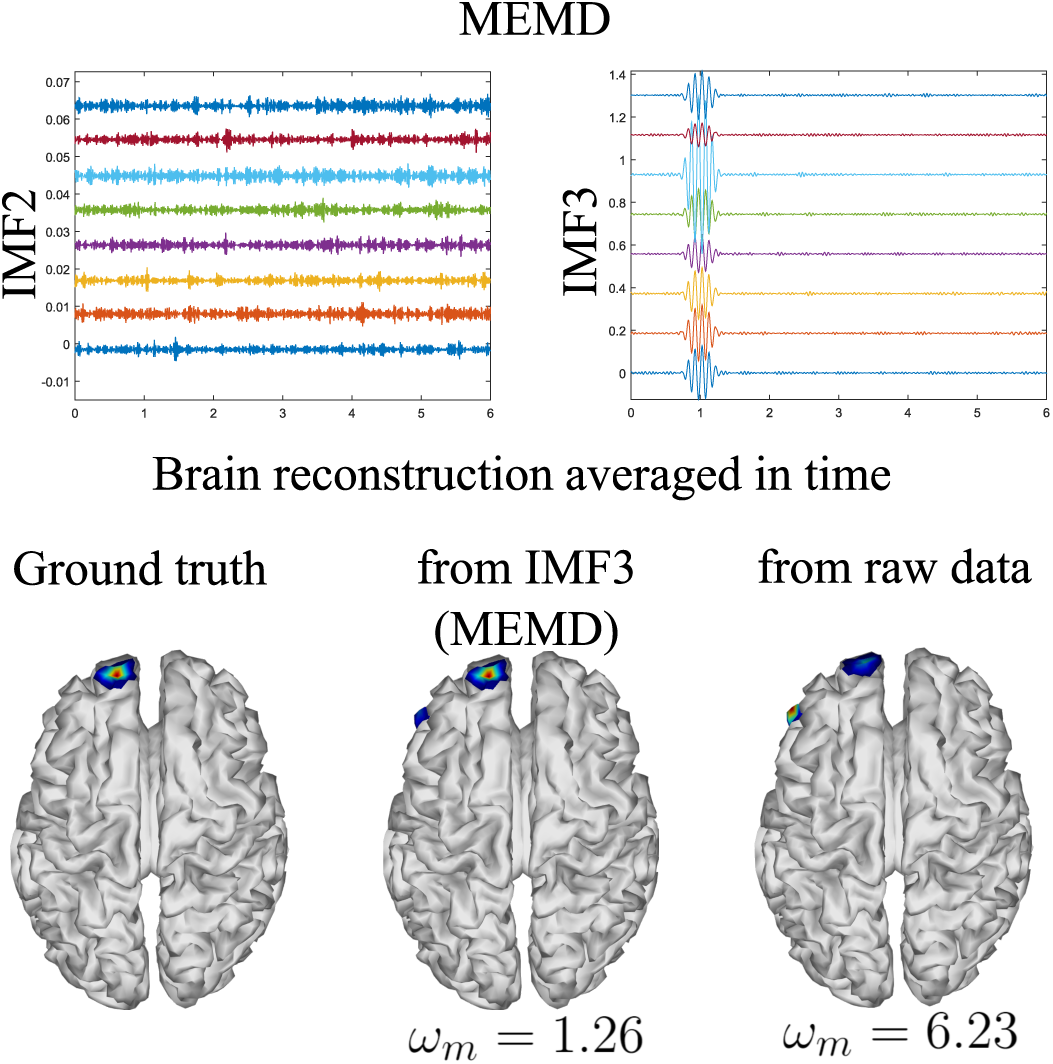
MEMD for one source with 10 Hz sinusoidal windowed activity and 8 EEG channels

As shown in Fig.5 for one source, the MEMD is able to unmix the frequency activity, this effect was also evident in the three and five active sources cases. Fig.6 depicts the main IMFs decomposed by MEMD over 16 EEG channels, and the resulting brain reconstruction for three sources case. It is noticeable that the decomposition by using the MEMD splits the activity clearly in three IMFs as follows: the activity around 20 Hz is shown in the IMF2, the activity around 12 Hz is shown in the IMF4, and the 4 Hz is shown in the IMF6. It is worth noting that there is no mode mixing in the MEMD decomposition. In addition, in terms of the Wasserstein metric, it can be seen that the MEMD-MSP achieves a *w*_*m*_ = 10.14, which is substantially less than raw-MSP value *w*_*m*_ = 15.57 when the neural activity reconstruction averaged over time is analyzed. Moreover, the MEMD-MSP reconstruction keeps the position of the three simulated sources, even if some spurious activity appears, in other hand, in the raw-MSP reconstruction, the second source located in left hemisphere visual cortex loss its position and spurious activity appears in different areas, even with more power than the first source, fact that is evident in the higher value of accuracy obtained.

**Fig. 6:**
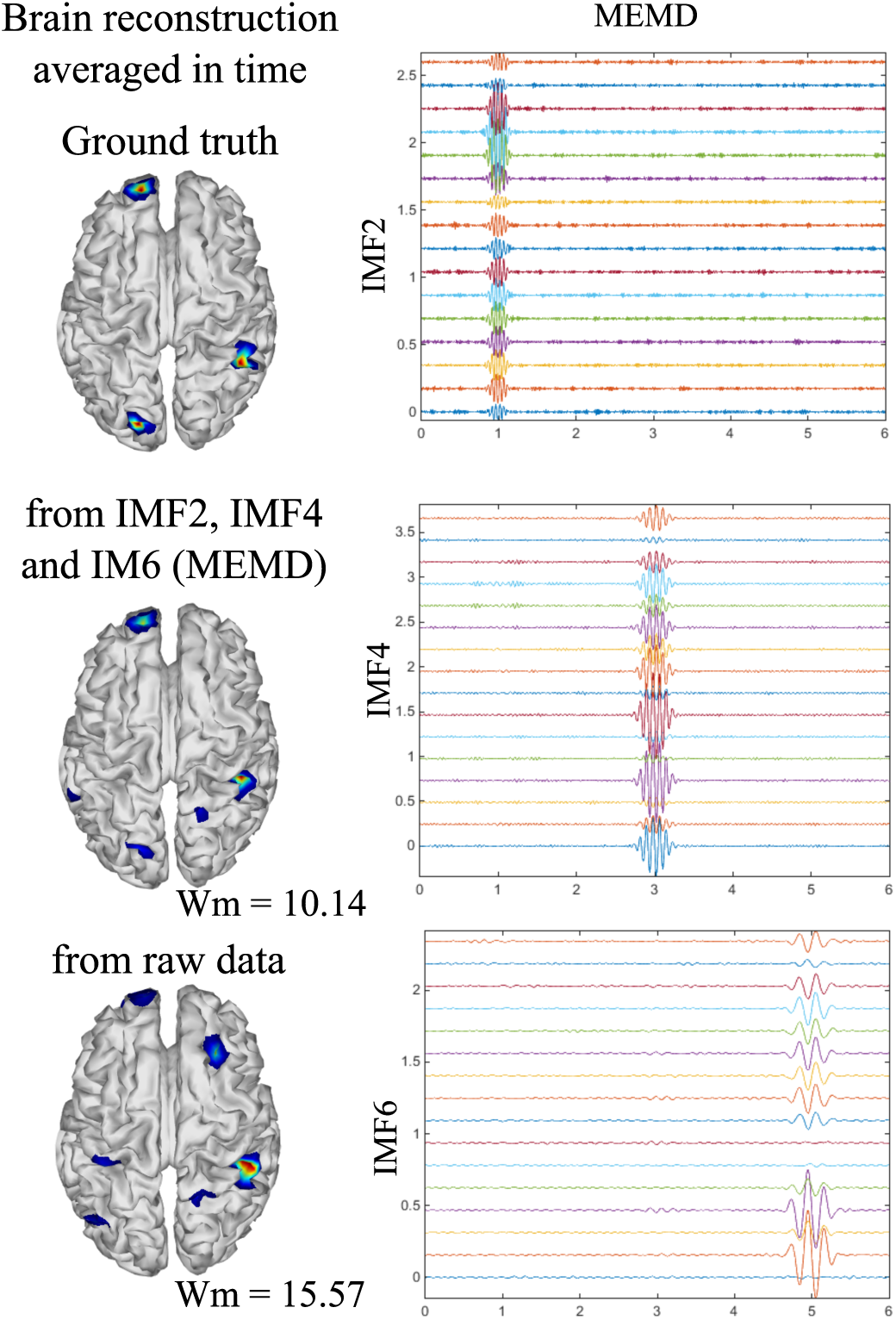
Ground truth activity, MEMD-MSP reconstruction, and raw-MSP reconstruction. The sources were simulated at 4, 12 and 20 Hz with a sinusoidal windowed activity and the source reconstruction was performed using 16 EEG channels. For the depicted MEMD-MSP reconstruction the IMFs 2, 4, and 6 (showed at right) were added to rebuild the EEG.

To analyze the three sources temporal reconstruction, the spatial and time evolution of the neural activity for the ground truth and the reconstructions by using MEMD-MSP and raw-MSP are shown in Fig. 7 for three time instants *t* = 1, *t* = 3 and *t* = 5 seconds with 16 EEG channels. It is clear that for each time instant the neural activity reconstruction obtained by the MEMD-MSP is better than the one obtained from the raw data in terms of the Wasserstein metric. The time evolution of the neural activity for the MEMD was obtained by mixing the resulting brain mapping for the IMF2, IMF4 and IMF6 of the MEMD, since the activity corresponding to each sources is clearly divided in the selected IMFs as shown in Fig. 6.

**Fig. 7:**
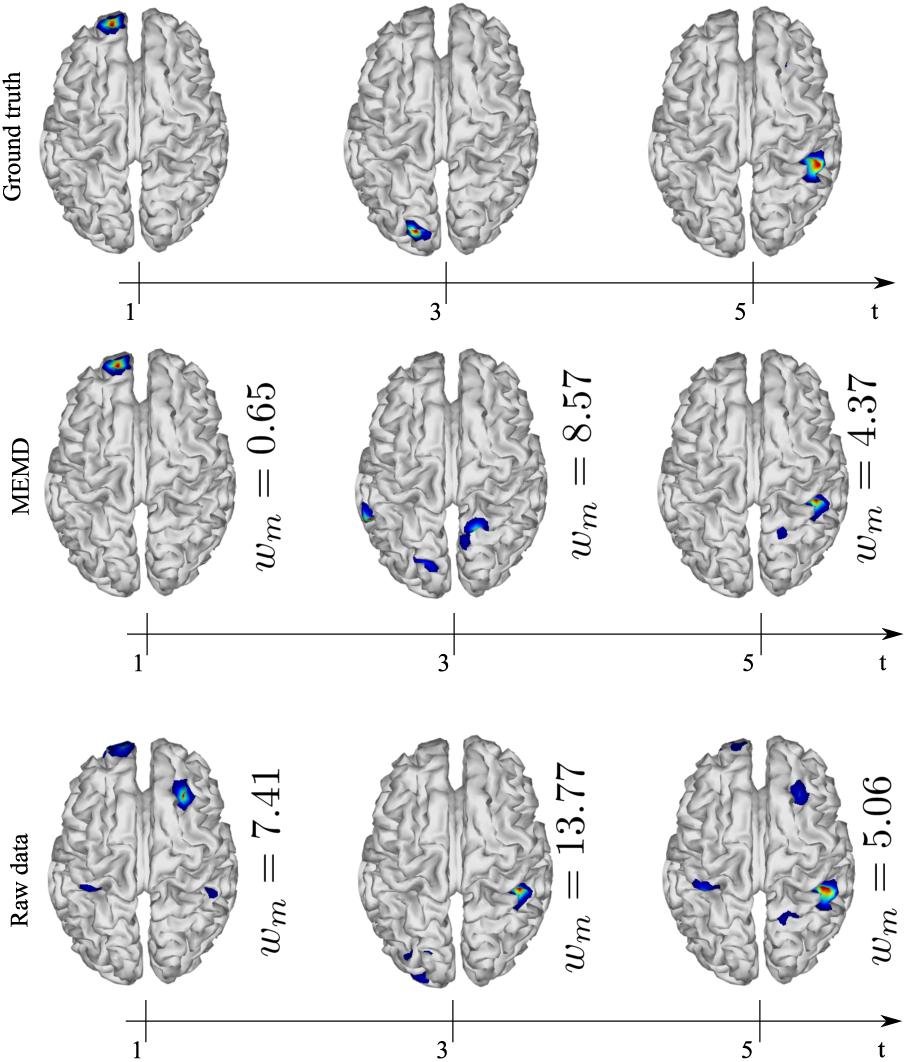
Ground truth, MEMD-MSP and raw-MSP neural activity mapping considering the evolution in time for three sources at time instants *t* = 1, *t* = 3 and *t* = 5 seconds with 16 EEG channels

A similar spatial and time evolution evaluation than showed in Fig. 7 is shown in Fig. 8, where the case of five active sources with 32 EEG channels was analyzed. In which the MEMD-MSP outperforms raw-MSP, the reconstructed neural activity with raw-MSP decrease the spatial accuracy in almost all the sources, just in the case of the third source at *t* = 3 the raw-MSP presents a low value than MEMD-MSP, however, the raw-MSP reconstruction contains several spurious activities at this third source, in addition, in the other sources reconstructions, it is possible to observe the effects of previous and posterior sources. In contrast, those effects are reduced when MEMD decomposition is applied, in which just in the fourth reconstructed source an attenuated activity from the third source could be seen, furthermore, the indexes of Wasserstein Metric are smaller than raw-MSP almost all the sources. This reduction of spurious activity and the attenuation of effects from other sources appear when the EEG signals were decomposed in IMF, and could be explained due to noisy information was rejected in the IMF selection process, and the mode mixing effects are attenuated by MEMD.

**Fig. 8:**
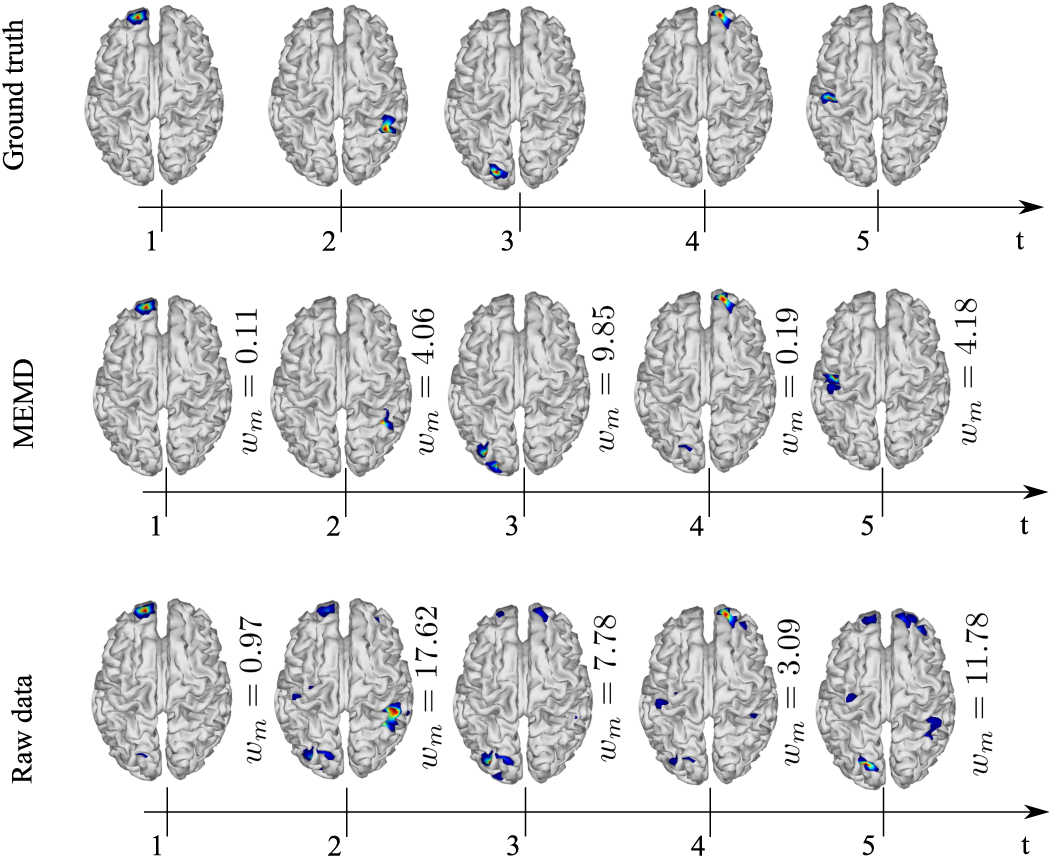
Ground truth, MEMD-MSP and raw-MSP neural activity mapping considering the evolution in time for five sources at time instants *t* = 1, *t* = 2, *t* = 3, *t* = 4 and *t* = 5 seconds with 32 EEG channels

Information of the reconstruction is shown in Fig. 9, where the mean and temporal reconstruction WM values are presented. In general, it could be seen the effects of the channels reduction, in which the source reconstruction with 8 channels increased the inaccuracy, specially when raw-MSP is applied. The results suggest that the brain reconstruction with MEMD-MSP can still be performed without loosing the accuracy by reducing the amount of EEG channels by a factor of two, in which, even if the WM index increase with the channel reduction, its slope is low and the reconstruction could be tolerated. In contrast, the raw-MSP method reconstruction presented a exponential increase when the electrode reduction is performed. In several cases, especially with less number of electrodes, the inclusion of the MEMD stage improves the performance of the brain mapping method significantly. Those significant differences presented in Fig. 9 were obtained by performing two-sided pairwise t-test with alpha level of *p <* 0.05 using IBM SPSS Statistics for Windows, version 24 (IBM Corp., Armonk, N.Y., USA), we compared the WM index between the MEMD-MSP and raw-MSP for mean reconstruction and for each source at time instants *t* = 1, *t* = 3, and *t* = 5 seconds with 32, 16, and 8 electrodes.

**Fig. 9:**
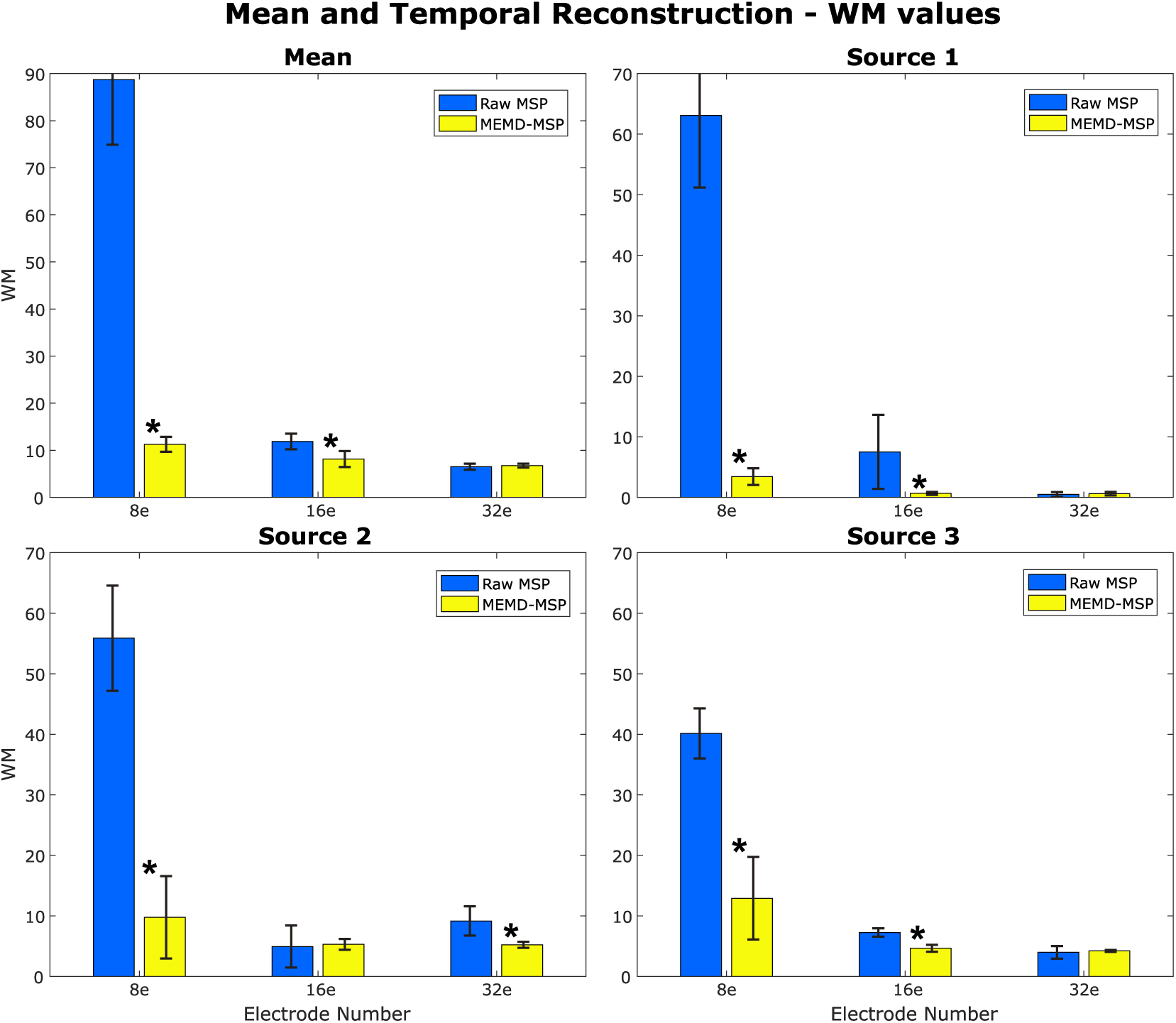
WM mean and standard deviation of the mean and temporal reconstruction with three number of electrodes, and considering three active sources in the brain at times 1s, 3s, and 5s.*Significant improvements with p*<*0.001

In general, the results over the synthetic EEG signals suggest that the use of the extracted information by MEMD improve the brain mapping method MSP. In all the analyzed cases, the MEMD-MSP has attenuated the appearance of spurious activity, and during the electrodes reduction, the joint of the methods keep the spatial accuracy.

### B. Results over EEG Dataset

We applied the methodology showed in Fig.3, the dataset was processed using with MEMD as a pre-processing step, the brain mapping solutions were obtained using MSP for MEMD and directly from the ERP data, and then, each subject average was compared with its own ground truth activity. A general vision of the results is shown in Fig. 10, where is presented the general mean of the WM and its standard deviation across all the subjects and conditions, comparing raw-MSP and MEMD-MSP with the ground truth activity by number of electrodes. It can be seen that the electrode reduction applied affect directly the quality of the source reconstruction, in which the solutions with MEMD-MSP have a lower mean and standard deviation that the raw-MSP in all the cases. The inaccuracy of raw-MSP increase as the electrodes are reduced with a high slope. In contrast, the MEMD-MSP keep a constant quality index when the brain mapping is performed with 32 and 16 electrodes, and increase when 8 electrodes were used, however, with 8 electrodes the MEMD-MSP reached a WM value and standard deviation similar to the obtained by raw-MSP with 32 electrodes, where no significant differences were found.

**Fig. 10:**
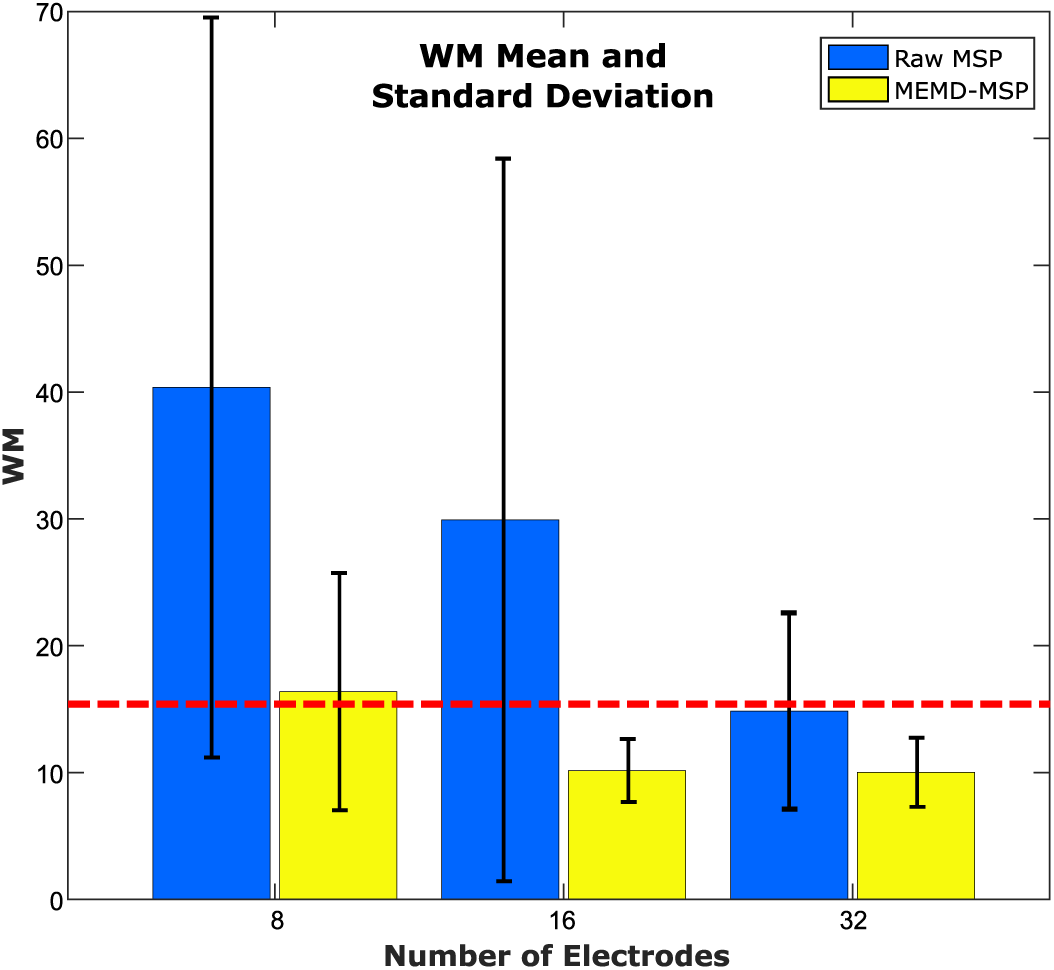
WM mean and standard deviation according to the number of electrodes

In order to provide an additional point of view of the results, the WM indexes were labeled according to the condition of the EEG signals (familiar or scrambled faces) and the quantity of electrodes used for performing the inverse solutions. The WM means according to the type of experiment and number of electrodes are shown in Fig. 11. In which the results show that for the case of scrambled faces with 8 electrodes with MEMD-MSP the WM mean is slightly than the obtained with raw-MSP with 32 electrodes, in other hand, the for the case of familiar faces the raw-MSP with 32 electrodes get a lower WM value than MEMD-MSP with 8 electrodes, however, when the comparison is between the same condition and the same number of electrodes, in all the cases, MEMD-MSP outperforms MSP.

**Fig. 11:**
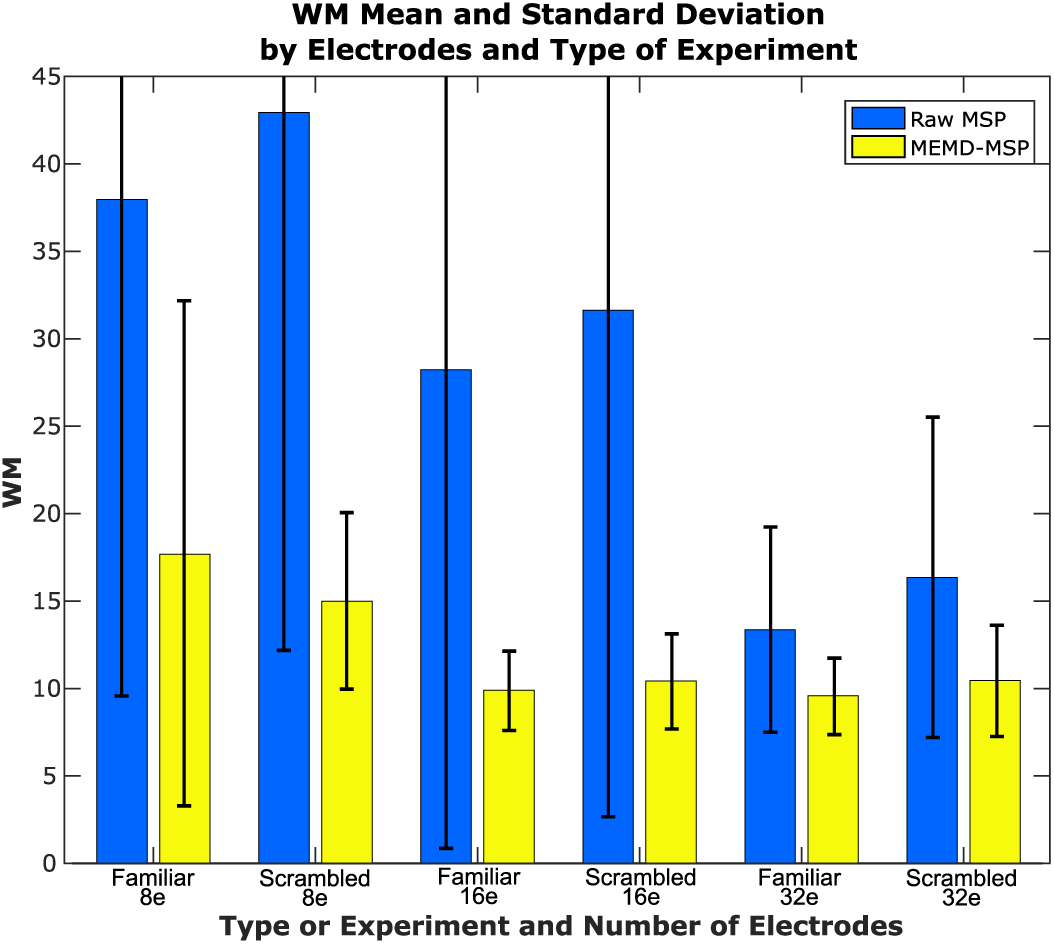
WM mean and standard deviation according to the type of experiment and number of electrodes

The improvement of the results applying the Multivariate EMD can be explained due to the separation of IMFs as is shown by Fig.12. Where is presented a 8 electrodes EEG signals and the IMF4 EEG reconstruction obtained with MEMD, the figure depicts how is extracted the main information of the evoked response around the established ROI, where the sparse temporal and frequency information provided by the IMF is enough to get a better source localization of the activity than using all the components of the EEG signal.

**Fig. 12:**
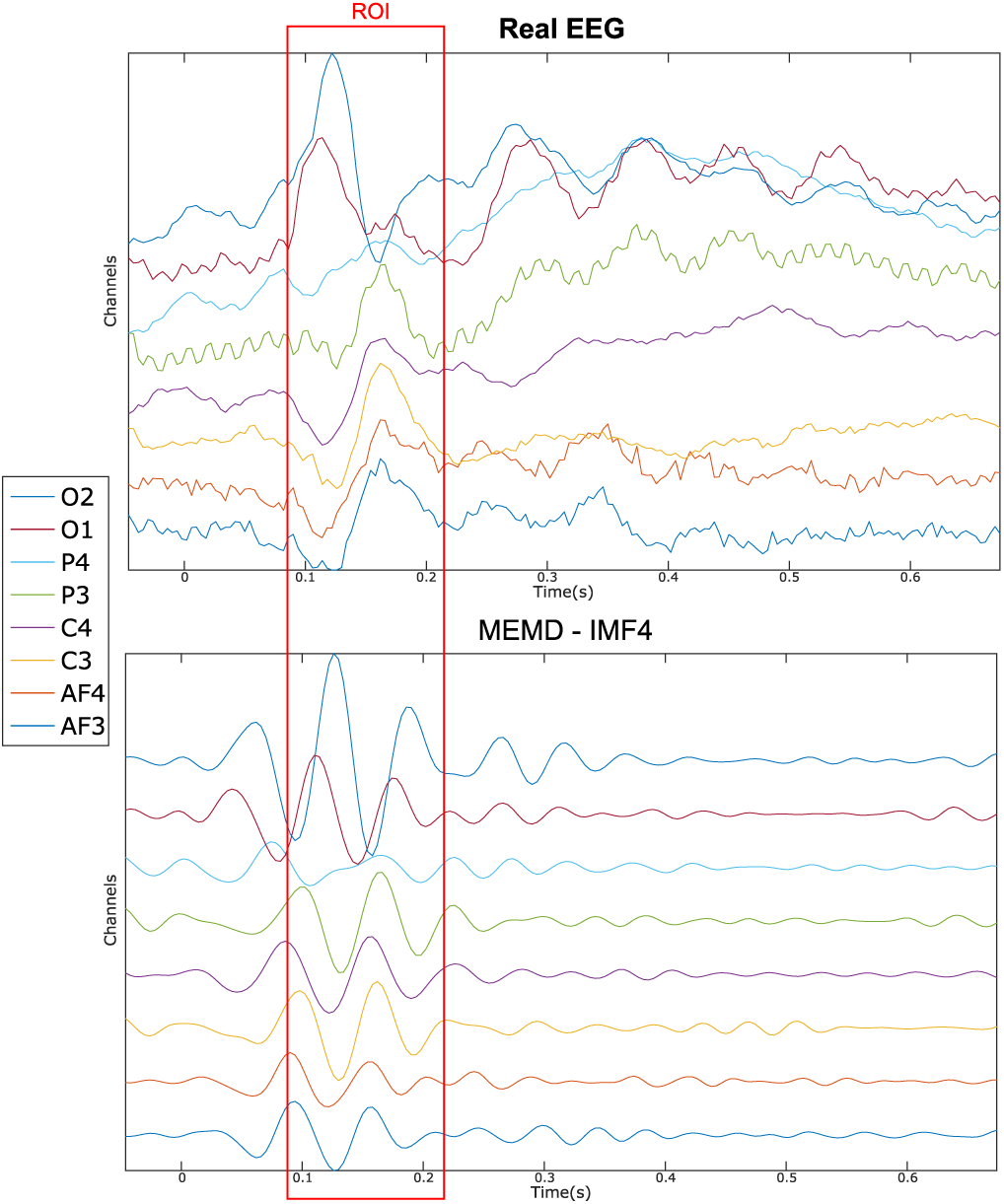
8 channel real EEG (Top) and the IMF4 decomposition (Botton)

Fig.13 shows the location of the neural activity in the brain, it activity has been found in the visual cortex, as the results presented in Wakeman and Henson (2015) and Henson et al. (2011) with a multi-modal technique involving EEG+MEG+fMRI. The figure provide the ground truth activity and the brain mapping reconstruction with 8, 16 and 32 electrodes with raw-MSP and MEMD-MSP.

**Fig. 13:**
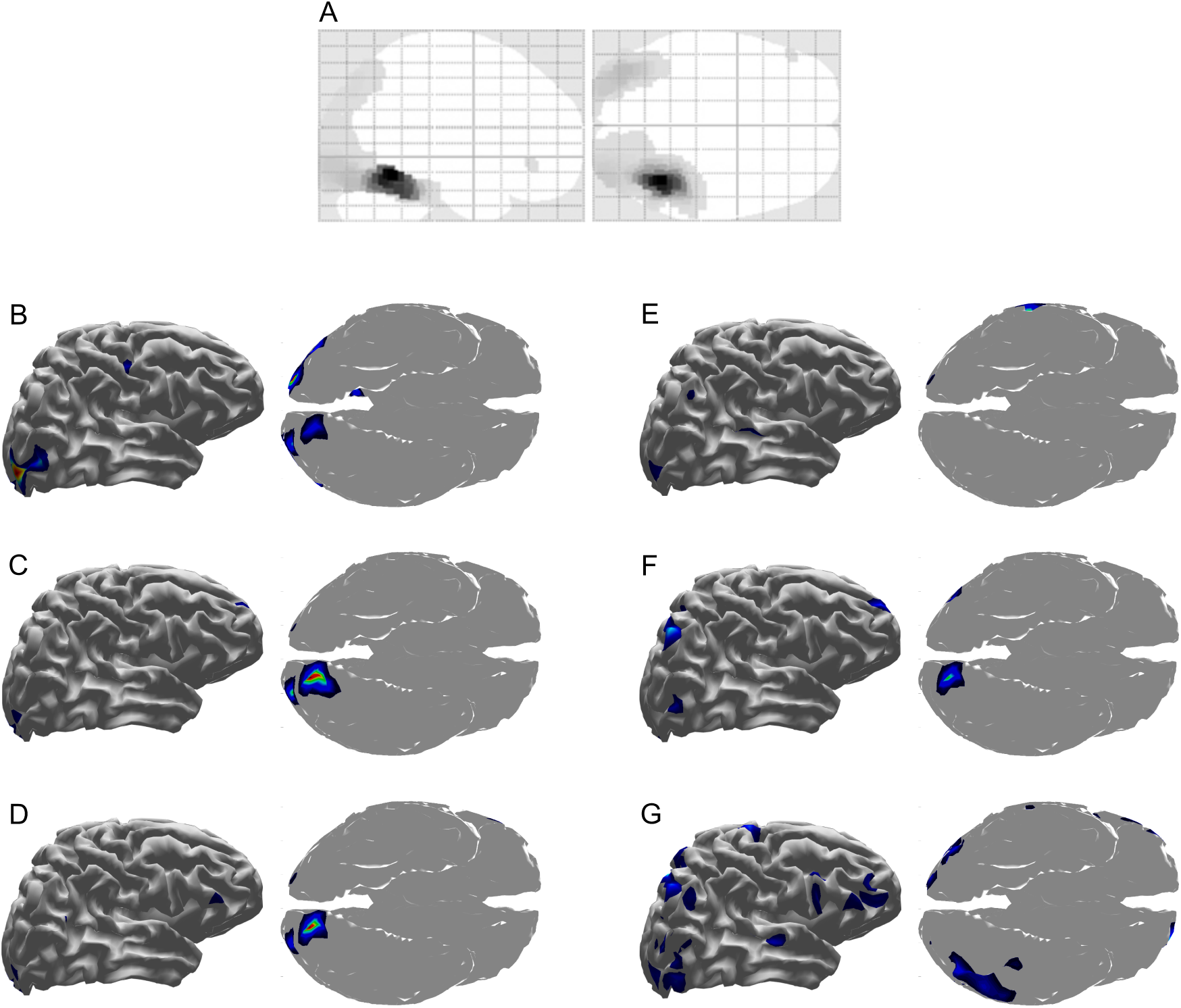
Brain activity reconstruction with a multi-modal technique involving EEG+MEG+fMRI ground truth activity from Henson et al. (2011) **(A)**, using MEMD-MSP with 8 electrodes **(B)**, 16 electrodes **(C)**, and 32 electrodes **(D)**, using raw-MSP with 8 electrodes **(E)**, 16 electrodes **(F)**, and 32 electrodes**(G)**.

From Fig.13, it can be seen that the reconstructions using MEMD-MSP shown a low variation across the diferent number of electrodes involved in the source localization, in the other hand, the without pre-processing the data, the raw-MSP reconstructions have a higher variation in localization and intensity of the source between electrodes, moreover, those solutions present activity in the diferent brain areas, when the MEMD method focused just in the visual cortex, which is directly involved during the visual face stimulus, therefore, the use of some IMFs provided by MEMD have resulted in a constant localization of the neural activity and an attenuation of background activity, which is represented by the lower values of WM.

## IV. Discussion and Conclusions

It is well known that the brain works under frequency rhythms between 0.5Hz at Delta rhythm to 45Hz in Gamma rhythm, the use of time-frequency decomposition methods to the EEG signals is generally applied to study the brain processes related to certain frequency activity and the changes in the brain waves oscillations during several experiments e.g. ERP studies. The EMD is a method that has shown the capability to separate signals using a time-frequency decomposition in different contexts, however the EEG signals are challenging due to the frequency proximity of the source activity, this condition makes that EMD solutions generally present mode mixing effects in the IMF decomposition, in which, the MEMD attenuate these effects when sources have a closer frequency as presented in Muñoz-Gutiérrez et al. (2018).

This study investigated the Multivariate version of EMD combined with source reconstruction algorithm MSP in order to evaluate the effects of MEMD as a pre-processing step during the calculation of the brain mapping solutions and their perform at several electrode montages with 32, 16 and 8 channels. We compared the solutions obtained with MEMD-MSP against raw-MSP for synthetic EEG signals, in which, we proposed three scenarios of source activity: one active source, three active sources, and five active sources, where the sources were simulated from 2 to 20Hz, and for real dataset of signals from a behavioral study of face perception as presented in Henson et al. (2003).

For all the evaluations performed, the experiments have shown that using a pre-processing step with MEMD improves the accuracy of the source reconstruction made by MSP. The results over synthetic and real EEG data show that the quality of the solutions with MEMD-MSP keep stable when a number of electrodes of 16 and 32 are used, and it just decreases using 8 channels. The reconstructions using MEMD-MSP with 8 channels achieved similar values than raw-MSP with 32 channels (shown in Fig.9 for simulated sources and in Fig.11 for real data), and significant differences were not found between those reconstructions, where is clear that adding the decomposition of MEMD and using selected information of IMF during the source reconstruction process make feasible to perform this process with low-density EEG. In which a low quantity of electrodes with sparse information of IMFs from MEMD can keep the accuracy of the source reconstruction and can reduce the effects of noise in the brain mapped solution.

We performed a temporal evaluation focused on ROIs defined by time windows around the appearance of sources in the synthetic EEG signals test and around the evoked activity for the real dataset. In general, the reconstruction with MEMD-MSP presented a clear attenuation of the background activity as can be shown in Fig. 6, 7, 8, and 13. This effect could be explained by the frequency decomposition and the mode mixing attenuation made by MEMD, in which the analysis of the frequency information of the EEG channels allows to MSP to focus on the source activity presented in the selected IMFs, as shown in Fig. 6, and 12, resulting in solutions with a lower index in terms of WM for the combination of methods.

In conclusion, the present study has shown that adding a priori time-frequency information as input to the source reconstruction method MSP, it is possible to find better solutions even when a sparse number of electrode’s information is used. That represents that with MEMD we could extract the main time-frequency information of the sources that are hidden over the electrodes data, and then used it in order to obtain a good quality reconstruction, comparable as the same MSP method with a high number of electrodes and without any prior information. Moreover, in the MEMD-MSP solutions, the source activity is clearly separable, producing an unmixing effect in the source space. The application of MEMD with other methods and the unmixed activity for brain connectivity will be studied in future, we consider that the presented method could be applied over Brain-Computer Interfaces applications and studies of brain connectivity due to the time-frequency information could be an important source of features at source space.

## Funding

This work was supported by the Norwegian University of Science and Technology NTNU, project “David and Goliath: single-channel EEG unravels its power through adaptive signal analysis”, also under the funding of COLCIENCIAS, research project 111077757982: Sistema de identificación de fuentes epileptogénicas basado en medidas de conectividad funcional usando registros electroencefalográficos e imágenes de resonancia magnética en pacientes con epilepsia refractaria: apoyo a la cirugía resectiva”.

## References

Boashash, B. (1992). Estimating and interpreting the instantaneous frequency of a signal. Part 1: Fundamentals. Proceedings of the IEEE, 80(4):520–538.

Bueno-Lopez, M., Giraldo, E., and Molinas, M. (2017a). Analysis of neural activity from EEG data based on EMD frequency bands. In 24th IEEE International Conference on Electronics, Circuits and Systems (ICECS), volume 1, pages 1–5, Batumi, Georgia. IEEE.

Bueno-Lopez, M., Molinas, M., and Kulia, G. (2017b). Understanding instantaneous frequency detection: A discussion of Hilbert-Huang Transform versus Wavelet Transform. In International Work-Conference on Time Series Analysis-ITISE, volume 1, pages 474–486, Granada, Spain. University of Granada.

Bueno-López, M., Muñoz-Gutiérrez, P. A., Giraldo, E., and Molinas, M. (2018). Analysis of epileptic activity based on brain mapping of eeg adaptive time-frequency decomposition. In Brain Informatics, pages 319–328, Cham. Springer International Publishing.

Bueno-López, M., Muñoz-Gutiérrez, P. A., Giraldo, E., and Molinas, M. (2019). Electroencephalographic source localization based on enhanced empirical mode decomposition. IAENG International Journal of Computer Science, 46(2):228–236.

Castano-Candamil, S., Hohne, J., Martínez-Vargas, J.-D., An, X.-W., Castellanos-Domínguez, G., and Haufe, S. (2015). Solving the eeg inverse problem based on space*-*time*-*frequency structured sparsity constraints. NeuroImage, 118:598–612.

Friston, K., Ashburner, J., Kiebel, S., Nichols, T., and Penny, W., editors (2007). Statistical Parametric Mapping. Academic Press, London.

Friston, K., Harrison, L., Daunizeau, J., Kiebel, S., Phillips, C., Trujillo-Barreto, N., Henson, R., Flandin, G., and Mattout, J. (2008). Multiple sparse priors for the m/eeg inverse problem. NeuroImage, 39(3):1104–1120.

Friston, K., Henson, R., Phillips, C., and Mattout, J. (2006). Bayesian estimation of evoked and induced responses. Human Brain Mapping, 27(9):722–735.

Fukushima, M., Yamashita, O., Knösche, T. R., and aki Sato, M. (2015). MEG source reconstruction based on identification of directed source interactions on whole-brain anatomical networks. NeuroImage, 105:408–427.

Gramfort, A., Strohmeier, D., Haueisen, J., Hamalainen, M., and Kowalski, M. (2013). Time-frequency mixed-norm estimates: Sparse m/eeg imaging with non-stationary source activations. NeuroImage, 70(0):410–422.

Hämäläinen, M. S. and Ilmoniemi, R. J. (1994). Interpreting magnetic fields of the brain: minimum norm estimates. Medical & Biological Engineering & Computing, 32(1):35–42.

Haufe, S., Nikulin, V. V., Ziehe, A., Müller, K.-R., and Nolte, G. (2008). Combining sparsity and rotational invariance in eeg/meg source reconstruction. NeuroImage, 42(2):726–738.

Henson, R. N., Ganel, T., and Otten, L. J. (2003). Electrophysiological and Haemodynamic Correlates of Face Perception, Recognition and Priming. Cereb. Cortex, 13:793–805.

Henson, R. N., Mattout, J., Phillips, C., and Friston, K. J. (2009). Selecting forward models for MEG source-reconstruction using model-evidence. NeuroImage, 46(1):168–176.

Henson, R. N., Mattout, J., Singh, K. D., Barnes, G. R., Hillebrand, A., and Friston, K. (2007). Population-level inferences for distributed MEG source localization under multiple constraints: Application to face-evoked fields. NeuroImage, 38(3):422–438.

Henson, R. N., Wakeman, D. G., Litvak, V., and Friston, K. J. (2011). A Parametric Empirical Bayesian Framework for the EEG/MEG Inverse Problem: Generative Models for Multi-Subject and Multi-Modal Integration. Frontiers in Human Neuroscience, 5(August):1–16.

Hou, T. Y. and Shi, Z. (2013). Data-driven time-frequency analysis. Applied and Computational Harmonic Analysis, 35(2):284–308.

Huang, N. E., Shen, Z., Long, S. R., Wu, M. C., Shih, H. H., Zheng, Q., Yen, N.-C., Tung, C. C., and Liu, H. H. (1998). The empirical mode decomposition and the Hilbert spectrum for nonlinear and non-stationary time series analysis. Proceedings of the Royal Society of London A: Mathematical, Physical and Engineering Sciences, 454(1971):903–995.

Jatoi, M. A. and Kamel, N. (2018). Brain source localization using reduced eeg sensors. Signal, Image and Video Processing, 12(8):1447–1454.

Khosropanah, P., Ramli, A. R., Lim, K. S., Marhaban, M. H., Ahmedov, A., and Jong-Mo, S. (2018). Fused multivariate empirical mode decomposition (memd) and inverse solution method for eeg source localization. Biomedical Engineering / Biomedizinische Technik, 63(4):467–479.

Lin, K. Y., Chen, D. Y., and Tsai, W. J. (2016). Face-based heart rate signal decomposition and evaluation using multiple linear regression. IEEE Sensors Journal, 16(5):1351–1360.

López, J., Litvak, V., Espinosa, J., Friston, K., and Barnes, G. (2014). Algorithmic procedures for bayesian meg/eeg source reconstruction in spm. NeuroImage, 84:476 –487.

Men-Tzung, L., Kun, H., Yanhui, L., Peng, C., and Vera, N. (2008). Multimodal pressure-flow analysis: Application of Hilbert Huang transform in cerebral blood flow regulation. EURASIP Journal on Advances in Signal Processing, 2008(1):1–15.

Muñoz-Gutiérrez, P. A., Giraldo, E., Bueno-López, M., and Molinas, M. (2018). Localization of active brain sources from eeg signals using empirical mode decomposition: A comparative study. Frontiers in Integrative Neuroscience, 12:55.

Muñoz-Gutiérrez, P. A., Giraldo, E., Bueno-López, M., and Molinas, M. (2019). Automatic selection of frequency bands for electroencephalographic source localization. In 2019 9th International IEEE/EMBS Conference on Neural Engineering (NER), pages 1179–1182.

Okcana, K. (2016). Definition of the instantaneous frequency of an electroencephalogram using the Hilbert transform. Advances in Bioscience and Bioengineering, 4(5):43–50.

Pascual-Marqui, R. D., Michel, C., and Lehmann, D. (1994). Low resolution electromagnetic tomography: a new method for localizing electrical activity in the brain. International Journal of Psychophysiology, 18(1):49–65.

Rehman, N. and Mandic, D. P. (2009). Multivariate Empirical Mode Decomposition. Proceedings of the Royal Society A, 466(1):1291–1302.

Rilling, G. and Flandrin, P. (2008). One or two frequencies? The empirical mode decomposition answers. IEEE Transactions on Signal Processing, 56(1):85–95.

Rubner, Y., Tomasi, C., and Guibas, L. J. (2000). The earth mover’s distance as a metric for image retrieval. International Journal of Computer Vision, 40(2):99–121.

She, Q.-s., Ma, Y.-l., Meng, M., Xi, X.-g., and Luo, Z.-z. (2017). Noise-assisted memd based relevant imfs identification and eeg classification. Journal of Central South University, 24(3):599–608.

Soleymani, M., Asghari-Esfeden, S., Fu, Y., and Pantic, M. (2016). Analysis of EEG signals and facial expressions for continuous emotion detection. IEEE Transactions on Affective Computing, 7(1):17–28.

Subha, D. P., Joseph, P. K., Acharya U R., and Lim, C. M. (2010). EEG signal analysis: A survey. Journal of Medical Systems, 34(2):195–212.

Wakeman, D. G. and Henson, R. N. (2015). A multi-subject, multi-modal human neuroimaging dataset. Scientific Data, 2:150001 EP –. Data Descriptor.

Wu, Z. and Huang, N. E. (2009). Ensemble Empirical Mode Decomposition: A noise-assisted data analysis method. Advances in Adaptive Data Analysis, 1(1):1–41.

Xu, D., Xia, Y., and Mandic, D. P. (2016). Optimization in quaternion dynamic systems: Gradient, hessian, and learning algorithms. IEEE Transactions on Neural Networks and Learning Systems, 27(2):249–261.

Xue, Y.-J., xing Cao, J., Du, H.-K., Zhang, G.-L., and Yao, Y. (2016). Does mode mixing matter in EMD-based highlight volume methods for hydrocarbon detection? experimental evidence. Journal of Applied Geophysics, 132(Supplement C):193–210.

Yin, Y., Cao, J., and Tanaka, T. (2012). Eeg energy analysis based on memd with ica pre-processing. In Proceedings of The 2012 Asia Pacific Signal and Information Processing Association Annual Summit and Conference, pages 1–5.

Zahra, A., Kanwal, N., ur Rehman, N., Ehsan, S., and McDonald-Maier, K. D. (2017). Seizure detection from eeg signals using multivariate empirical mode decomposition. Computers in Biology and Medicine, 88:132 –141.

